# Copper imbalance linked to oxidative stress and cell death during Zika virus infection in human astrocytes

**DOI:** 10.1101/2021.12.29.474370

**Authors:** Teresa Puig-Pijuan, Leticia R. Q. Souza, Carolina da S. G. Pedrosa, Luiza M. Higa, Fabio Luis Monteiro, Amilcar Tanuri, Rafael H. F. Valverde, Marcelo Einicker-Lamas, Stevens Kastrup Rehen

**Affiliations:** D’Or Institute for Research and Education (IDOR), Diniz Cordeiro 30, 22281-100, Rio de Janeiro, Brazil; Carlos Chagas Filho Biophysics Institute, Federal University of Rio de Janeiro, Av Carlos Chagas Filho 373, 21941-902, Rio de Janeiro, Brazil; Department of Genetics, Institute of Biology, Federal University of Rio de Janeiro, Rodolpho Paulo Rocco SN, 21941-617, Rio de Janeiro, Brazil

**Keywords:** copper, Zika Virus, oxidative stress, human astrocytes, viral infection, neural cells

## Abstract

The Zika virus (ZIKV) caused neurological abnormalities in more than 3500 Brazilian newborns between 2015 and 2020. Data have pointed to oxidative stress in astrocytes as well as to dysregulations in neural cell proliferation and cell cycle as important events accounting for the cell death and neurological complications observed in Congenital Zika Syndrome. Copper imbalance has been shown to induce similar alterations in other pathologies, and disturbances in copper homeostasis have already been described in viral infections. For this reason, we investigated copper homeostasis imbalance as a factor that could contribute to the cytotoxic effects of ZIKV infection in iPSC-derived human astrocytes. Our results show that ZIKV infection leads to a downregulation of one of the transporters mediating copper release, ATP7B protein. We also observed the activation of mechanisms that counteract high copper levels, including the synthesis of copper chaperones and the reduction of the copper importer protein CTR1. Finally, we show that chelator-mediated copper sequestration in ZIKV-infected astrocytes reduces the levels of reactive oxygen species and improves cell viability, but does not change the overall percentage of infected cells. In summary, our results show that copper homeostasis imbalance plays a role in the pathology of ZIKV in astrocytes, indicating that it may also be a factor accounting for the developmental abnormalities in the central nervous system following viral infection. Evaluating micronutrient levels and the use of copper chelators in pregnant women susceptible to ZIKV infection may be promising strategies to manage novel cases of congenital ZIKV syndrome.

## Introduction

Zika virus (ZIKV) is an enveloped single-stranded, positive-sense RNA virus that belongs to the *Flaviviridae* family and genus *Flavivirus*. Although ZIKV has been known since 1947 [1], it only became a major concern in 2015, after the correlation with severe neurodevelopmental abnormalities in newborns whose mothers were infected during pregnancy. ZIKV-related birth complications, which include microcephaly, brain parenchyma calcifications, congenital contractures and hypertonia, were collectively defined as Congenital Zika Syndrome (CZS). From 2015 to 2019, CZS was confirmed in more than 3000 newborns and stillbirths only in Brazil [2]. Although CZS occurrence dropped after 2016, cases of ZIKV infection are still being reported every year [3].

The devastating effects of the virus in newborn children initiated a global research effort to understand the mechanism of ZIKV pathogenicity. It was soon discovered that the virus was able to infect and cause proliferative defects and cell death in neural progenitor cells, which might account for brain growth impairments [4], [5]. Other cell types were also susceptible to infection including oligodendrocytes, astrocytes, microglia, endothelial cells from the blood-brain barrier (BBB), and neurons [6]. Of these cell types, astrocytes represent the most abundant cells in the brain and are responsible for a number of neurodevelopmental functions including axonal guidance, synaptogenesis, and neuronal survival, which implies that any impairment on their function can severely impact brain development [7]. In addition, it has been suggested that astrocytes might be the first site of ZIKV infection in the brain [8] and that they facilitate viral spread in the CNS, since they are highly susceptible to ZIKV infection and the virus can efficiently replicate inside them [9].

Astrocytes are also crucial for other functions in the CNS, including the regulation of copper homeostasis [10]. Copper is an essential metal which is important in many bodily functions due to its role as a cofactor of several key enzymes, but also due to its essentiality in signaling pathways like MEK/ERK, where copper is needed for ERK1/2 phosphorylation [11], [12]. In addition, copper can display both prooxidant and antioxidant effects since it works as a cofactor for superoxide dismutase 1 (SOD1) but, in excess, it can also promote the generation of reactive oxygen species (ROS) [13]. Copper imbalance can also impair mitochondrial function due to its role as a cofactor for cytochrome c oxidase [14], [15]. Because of its biological importance and its potential damaging effects, copper metabolism is tightly regulated in most cells, including astrocytes, by a set of transporters and chaperones that control copper internalization, distribution and secretion [16].

These include CTR1, the main copper internalization protein; ATP7A and ATP7B, which are each expressed in different tissues and mediate both copper excretion and delivery to copper-containing proteins in the Golgi; copper chaperones COX17, ATOX1 and CCS, which are responsible for binding free copper and delivering it to the mitochondria, copper ATPases and superoxide dismutase I, respectively; and glutathione and metallothioneins which capture free copper to avoid oxidative damage. The central nervous system (CNS) is especially sensitive to any perturbation in copper homeostasis, as evidenced by neurological function impairment in diseases caused by mutations in copper ATPases genes, and by the fact that copper dysmetabolism has been reported in neurodegenerative diseases including Parkinson’s and Alzheimer’s diseases [17], [18]. Interestingly, many of the deleterious effects of ZIKV in neural cells, including cell cycle alterations, oxidative stress, mitochondrial failure and inflammation [19], [20] are also associated with copper imbalance [21]. Previous studies have shown that intracellular copper balance is required for influenza A virus replication [22], while other viral infections cause important alterations in copper levels [23], [24]. Here, we hypothesized that disturbance of copper homeostasis is a mechanism of ZIKV to promote oxidative stress and cell death in astrocytes. Our findings offer a new perspective on the importance of exploring metal homeostasis during brain development and viral infections, directing for new therapeutic approaches to prevent CZS.

## Methods

### Cell cultures

Human induced pluripotent stem cells (hiPSC) were obtained by differentiation of skin fibroblasts from two healthy subjects. Neural stem cells (NSC) were differentiated from hiPSC according to the protocol described by Yan et al. [25]. Human astrocytes were differentiated from NSCs using the protocol described in Trindade et al. [26]. The hiPSC lines used were previously described in Casas et al. [27]; and the hiPSC derived-astrocytes were characterized in Ledur et al. [19]. Reprogramming of human cells was approved by the ethics committee of Copa D’Or Hospital (CAAE number 60944916.5.0000.5249, approval number 1.791.182). Human cell experiments were performed according to Copa D’Or Hospital regulations.

### ZIKV production and infection

ZIKV was isolated from the serum of a patient from the state of Pernambuco (PE) (ZIKV PE243/2015, GenBank accession number: KX197192.1, Recife /Brazil). ZIKV was propagated and tittered as described in Ledur et al. [19]. For the infection experiments, cells were washed once with PBS and then incubated for 2 hours with the inoculum containing ZIKV at a multiplicity of infection (MOI) of 1, in serum-free medium. Infection control was performed by incubating cells with an equivalent volume of conditioned supernatant from uninfected C6/36 cells (mock). After the incubation period, the inoculum was removed and cells were returned to fresh culture medium. For some experiments, cells were treated with either 200 μM bathocuproinedisulfonic acid (BCS) or 100 μM CuCl_2_ immediately after the inoculum incubation period.

### ZIKV titration by plaque assay

To address the production of ZIKV infectious virus particles after treatment, plaque assay was performed using Vero CCL-81™ (ATCC) cells. Ten-fold serial dilutions of conditioned medium from infected/treated cells were inoculated into confluent Vero monolayers. After 1 h of incubation at 37 °C and 5% CO_2_, the inoculum was removed and cells were overlaid with semisolid medium constituted of alpha-MEM (Gibco) containing 1.25% carboxymethyl cellulose (Sigma Aldrich), 1% fetal bovine serum (Gibco) and 1% penicillin-streptomycin. Cells were further incubated for 5 days and then fixed with 4% formaldehyde. Cell monolayers were stained with 0.5% crystal violet in 20% ethanol for plaque visualization. Titers were expressed as plaque forming units (PFU) per milliliter.

### Gene expression analysis

RNA was isolated from frozen cell pellets using *Pure Link RNA Mini kit* (Invitrogen) according to manufacturer instructions. RNA samples were treated with DNase I (Invitrogen) and cDNA was synthesized using M-MLV reverse transcriptase kit (Invitrogen) according to manufacturer instructions. Conventional PCR followed by 2% agarose gel electrophoresis was performed to detect ZIKV entry receptors. For gene expression analysis, qPCR was performed using Fast Plus EvaGreen^®^ qPCR master mix High ROX (Biotium). Primers for genes of interest and genes of reference (GAPDH and HPRT1) are listed in Supplementary Table 1.

### Western blot

For sample preparation, cells were trypsinized and lysed using a lysis buffer containing 50 mM potassium phosphate (pH 7.8), 0.1 mM EDTA, 0.1% Triton, 7% PMSF and 0.5% aprotinin. Samples were vortexed 3 times for 10 seconds and manually homogenized using a Potter-Elvehjem homogenizer. SDS-PAGE electrophoresis was performed using 50 μg protein and samples were then transferred to nitrocellulose membranes for 1h and 30 min at 350 mA. Membranes were blocked with 5% nonfat dry milk for 1h at room temperature. Primary antibodies against CTR1 (sc-66847, Santa Cruz Biotechnology) and β-actin (A5441, Sigma Aldrich) were diluted 1:1,000 in Tris-buffered saline (TBS) and incubated for 2h at room temperature. Membranes were washed three times with TBS containing 0.1% Tween 20. Secondary antibodies were diluted 1:5,000 in TBS and incubated for 1h at room temperature in a shaker. Protein bands were visualized using Immobilon Forte Western HRP Substrate (Millipore) and ChemiDoc system (BioRad).

### Immunostaining

For immunofluorescence, cells were plated in 24-well plates with coverslips. 72 hours after infection, cells were washed with PBS once and fixed with 4% paraformaldehyde for 20 minutes. Fixed cells were permeabilized with 0.1% Triton for 15 minutes and subsequently blocked for 2h with either 5% normal goat serum (for NS1 staining) or 3% bovine serum albumin (other markers) in PBS with 0.1% Triton. Samples were then incubated overnight at 4ºC with primary antibodies against non-structural protein from ZIKV (NS1) 1:500 (BF1225-06, BioFront Technologies) combined with Vimentin 1:1,000 (ab92547, Abcam). Cells were then washed and incubated with secondary antibodies 1:400 (A31573, Thermo Fisher Scientific; A11078, Invitrogen) for 2h at room temperature. Finally, coverslips were incubated with 300nM DAPI for 5 minutes, washed and mounted on glass slides with Aqua-Poly/Mount. Images from 9 randomly selected fields per coverslip were acquired using a Leica SP8 confocal microscope equipped with a 20x objective. Quantification of percentage of infected cells was done using ImageJ software.

### ROS analysis

For the detection of ROS, cells were plated in black-walled clear-bottom 96-well plates. Cells were incubated with either 5 μM DHE probe (D11347, Thermo Fisher Scientific) or 5 μM MitoSOX Red (M36008, Thermo Fisher Scientific) combined with 150nM MitoTracker Green (M7514, Thermo Fisher Scientific) and 1 μM Hoechst 33342 (H1399, Thermo Fisher Scientific) for 30 min at 37 ºC and 5% CO_2_. Images were acquired, maintaining cells at culture conditions (37 ºC and 5% CO_2_) using high-content imaging system Operetta^®^ (PerkinElmer) and analyzed using Harmony^®^ software.

### Cell viability

Astrocytes plated in 96-well transparent plates were washed once and incubated with 25 μg/mL Neutral Red in DMEM-F12 supplemented with 5% FBS and 1% penicillin-streptomycin for 3 hours at 37ºC and 5% CO_2_. After staining absorption, cells were washed once and then the dye was eluted with 50% ethanol/1% acetic acid solution for 20 min in a shaker. Absorbance was read at 540 nm using TECAN Infinite M200 PRO spectrophotometer. Cell viability values were normalized to untreated control absorbance values (set as 100%).

### Statistical analysis

For two-group comparisons, paired two-tailed Student’s t-test was performed. For three or more groups, we used one-way ANOVA followed by Tukey’s post hoc test. Data comprehend descriptive statistics expressed as mean ± SEM. P values are specified at each figure and at their respective legend, where *p < 0.05, **p < 0.01, ***p < 0.001, ****p < 0.0001.

## Results

### ZIKV causes alterations in gene expression and protein levels of copper regulatory proteins

Human iPSC-derived astrocytes are highly susceptible and permissive to ZIKV infection. In order to understand how copper metabolism disturbances might be involved in those cytopathic effects, we first analyzed the gene expression of the main proteins that regulate copper homeostasis in astrocytes infected with ZIKV (Figure 1a). The expression of ZIKV entry receptors in these astrocytes was confirmed by conventional PCR (Supplementary Figure 1).

**Figure 1.**
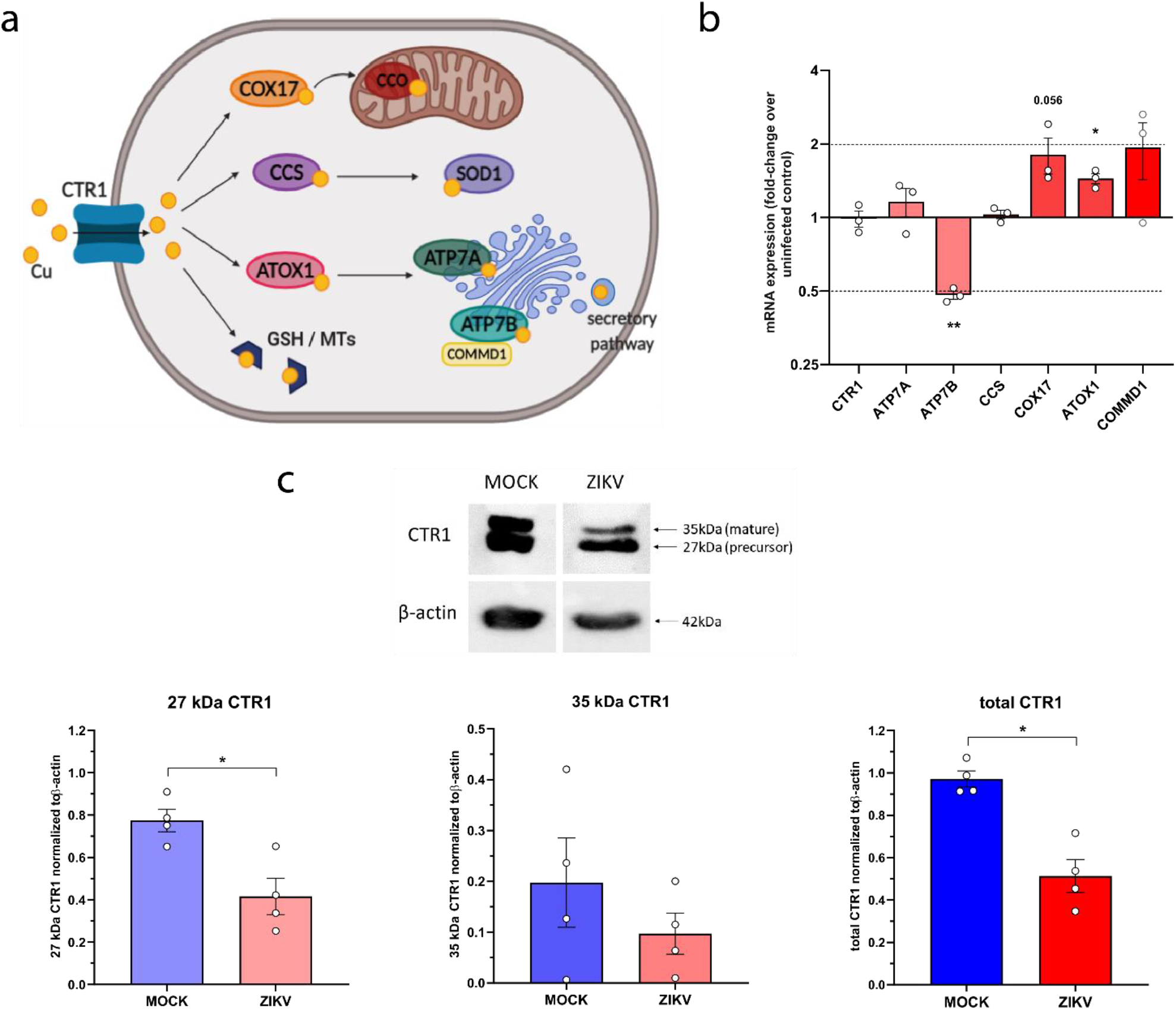
ZIKV causes alterations in gene expression and protein expression of copper regulatory proteins. a) Copper distribution pathways in mammalian cells. b) qPCR analysis of the main copper regulatory proteins in ZIKV-infected astrocytes compared to non-infected astrocytes, 48h post-infection. Results are expressed as fold-change over mock-infected controls. n=3. c) Western blot analysis of CTR1 in ZIKV-infected astrocytes 48h post-infection. Detection of CTR1 showed two bands corresponding to the precursor (27kDa) and mature (35kDa) forms of the protein. Protein levels were quantified by densitometry and normalized to β-actin levels. n=4. * p < 0.05; ** p < 0.01

Gene expression analysis performed 48h after infection showed a 0.48 ± 0.02 -fold significant reduction of ATP7B (p = 0.0026) (Figure 1b), which is one of the proteins responsible for copper release and delivery to cuproenzymes in the Golgi. ATOX1, a chaperone that transfers cytoplasmic copper to ATPases in the Golgi, had its gene expression significantly upregulated (1.44 ± 0.07 -fold, p = 0.0173). COX17, a chaperone that transfers copper from the cytoplasm to mitochondria, and COMMD1, which regulated ATPases localization and degradation, were also upregulated but not significantly (1.81 ± 0.30 -fold, p = 0.07 and 1.93 ± 0.51 -fold, p = 0.211, respectively).

Some of the main proteins that control copper homeostasis in the cell are mainly post-translationally regulated by degradation and translocation [28]. Copper internalizing transporter CTR1, for example, is usually located in the membrane when copper is required, and it is endocytosed and degraded when copper levels are high. Thus, we next performed Western Blot analysis to assess CTR1 protein levels in astrocytes during ZIKV infection. Densitometric analysis of CTR1 bands showed a significant reduction in the 27 kDa precursor form (0.77 ± 0.05 vs 0.42 ± 0.09, p = 0.025) and a tendency to reduction of the 35 kDa mature form of CTR1 (0.20 ± 0.09 vs. 0.10 ± 0.04, p = 0.430) in infected cells compared to non-infected cells (Figure 1c), which translated in a significant reduction of total CTR1 (0.97 ± 0.04, p = 0.026).

### Copper modulation reduces oxidative stress and cytotoxicity induced by ZIKV infection, but does not cause alterations in ZIKV infectivity

As seen in our previous work, oxidative stress contributes to the deleterious effect of ZIKV infection in astrocytes [19]. Copper imbalance can induce oxidative damage, therefore, the alterations in the mechanisms of copper homeostasis described here could contribute to oxidative stress in ZIKV-infected cells. To verify this, astrocytes were treated with 200 μM extracellular copper chelator BCS or 100 μM CuCl_2_ after incubation with the virus, and cytosolic and mitochondrial ROS levels were analyzed 48 hours after, using fluorescent probes DHE and MitoSOX Red respectively (Figure 2a,b). BCS and CuCl_2_ non-toxic concentrations were chosen after plotting cytotoxicity curves for each compound (Supplementary Figure 2). Treatment with BCS or CuCl_2_ did not significantly change ROS levels in uninfected cells (DHE: 4.94 ± 0.69 % DHE-positive cells in vehicle group vs 3.31 ± 0.97 and 3.51 ± 1.41 % positive cells in BCS-treated and CuCl_2_-treated cells, respectively; MitoSox: 5.63 ± 0.89 % MitoSox-positive cells in vehicle group vs 2.06 ± 0.77 % and 5.01 ± 1.07 % in BCS-treated and CuCl_2_-treated cells, respectively). As previously described, there was a significant increase in cytosolic and mitochondrial ROS in infected cells compared to uninfected cells (55.90 ± 3.27 % DHE-positive cells and 43.43 ± 2.32 % MitoSox-positive cells). BCS treatment in infected astrocytes significantly reduced ROS levels (33.57 ± 4.16 % DHE-positive cells and 22.99 ± 5.58 % MitoSox-positive cells), while treatment with CuCl_2_ did not significantly change ROS detection in infected astrocytes (45.19 ± 6.13 % DHE-positive cells and 36.87 ± 3.41 % MitoSox-positive cells). Thus, copper chelation is effective at reducing ROS levels in the cytoplasm and mitochondria of ZIKV-infected astrocytes.

**Figure 2.**
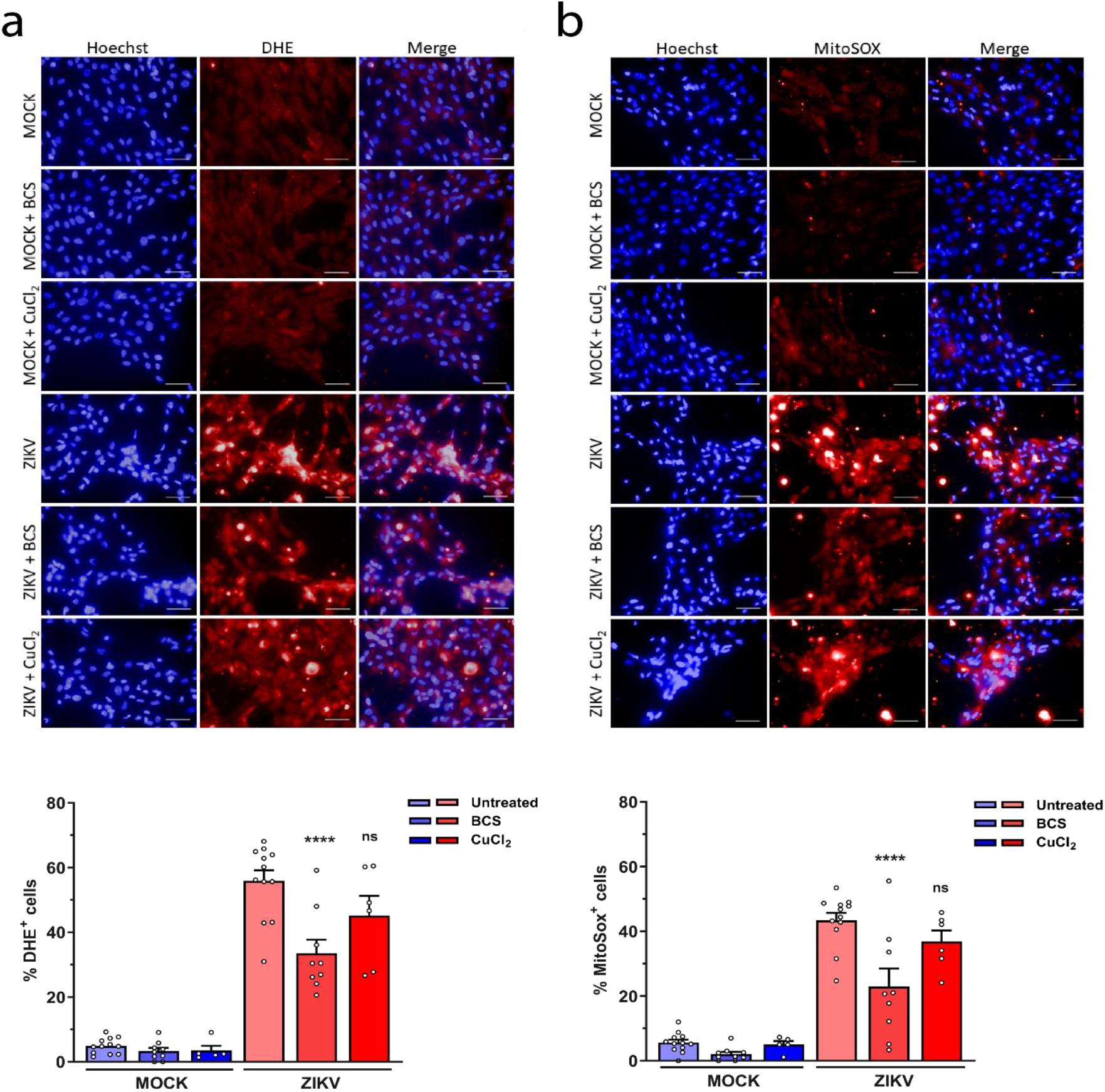
Effect of copper modulation on ZIKV-induced oxidative stress. Cytosolic (a) and mitochondrial (b) ROS levels were detected in infected (ZIKV) or uninfected (MOCK) human astrocytes treated with 200 μM of BCS or 100 μM CuCl_2_ using the ROS sensing probes DHE and MitoSOX Red, respectively. Nuclei were stained with Hoechst 33342. Quantification of ROS levels was expressed as percentage of DHE/MitoSOX positive cells over the total number of cells. Scale bar: 50 μm; n = 3. Data are shown as mean ± SEM; ns = not significant, ****p < 0.0001.

Oxidative stress is a possible inducer of cell death in ZIKV-infected cells and, as shown, copper chelation is able to reduce the amount of ROS produced during infection. Therefore, we assessed whether the modulation of copper levels would also be effective in reducing cell death caused by ZIKV infection. Cell viability was analyzed 72 hours after infection using neutral red uptake assay (Figure 3a). ZIKV infection caused a severe decrease in cell viability (5.66 ± 1.26 % viable cells), and treatment with BCS significantly improved viability (28.52 ± 3.60 % viable cells). Although treatment with CuCl_2_ in ZIKV-infected cells caused morphologic changes in astrocytes, there was a tendency to cell viability increase, but it was not significant (16.13 ± 0.80 % viable cells). In summary, the reduction of ROS levels and the improvement in cell viability after copper chelation indicate that changes in intracellular copper levels might be partly responsible for the oxidative damage that leads to cell death during ZIKV infection.

**Figure 3.**
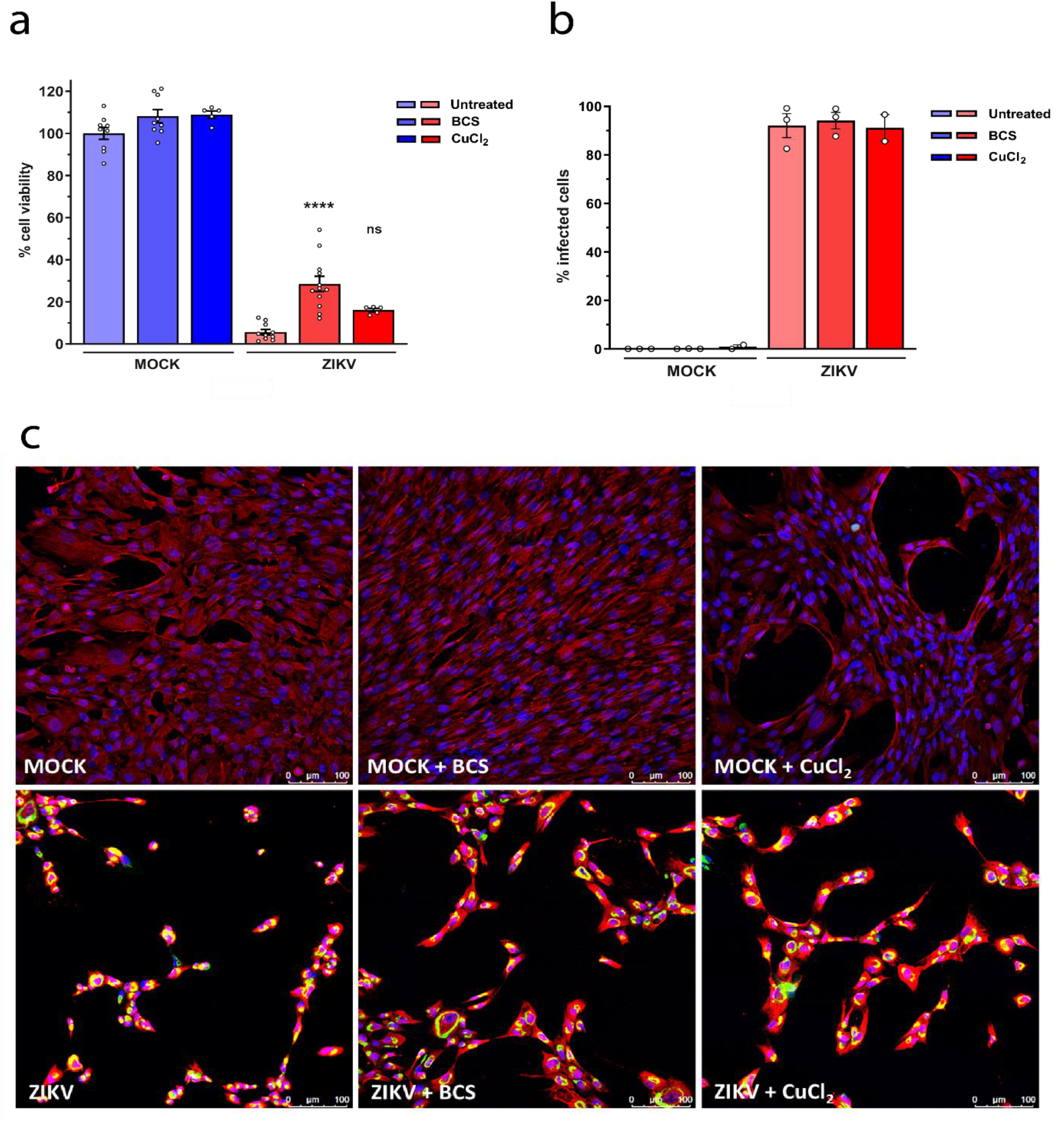
Effect of copper modulation on ZIKV cytotoxicity and infectivity. a) Cell viability in astrocytes infected with ZIKV and treated with CuCl_2_ or BCS. Viability was analyzed from the incorporation of neutral red dye and expressed as a percentage relative to the control group (untreated mock-infected cells). n = 3. b) Percentage of cells infected with ZIKV in astrocytes treated with copper (CuCl_2_) or copper chelator (BCS) 48h after infection. Data expressed as percentage of infected cells (NS1^+^ cells) from the total number of nuclei in each slide. c) Representative immunofluorescence of astrocytes stained with antibodies against the ZIKV NS1 protein (green) and vimentin (red). DAPI was used for nuclei staining (blue). Scale bar: 100 μm. n=3. Data are shown as mean ± SEM; ns = not significant, ****p < 0.0001.

Copper modulation might also affect viral replication or infectivity, as reported in other viral infections, which could also be related to the improvement of cell viability. We tested this hypothesis by evaluating the percentage of infected cells after copper modulation. For this, astrocytes were infected with ZIKV and treated with CuCl_2_ or BCS immediately after infection, and immunofluorescence for the NS1 viral protein was performed 48 hours after infection. Treatment with either BCS or CuCl_2_ did not significantly change the percentage of infected cells, which remained around 100% (Figure 3b,c). Plaque assay with conditioned medium also showed no changes in ZIKV infectivity after BCS or CuCl_2_ treatment (Supplementary Figure 3). Thus, the mechanisms that led to the improvement of cell viability during BCS treatment in infected astrocytes might not involve a reduction of infection levels.

## Discussion

Zika virus caused great commotion worldwide due to the congenital malformations caused to newborns during the 2015-2016 pandemic. Although many studies linked cell death and impaired proliferation of neural progenitors to microcephaly, it was soon noticed that astrocytes were also vulnerable to ZIKV infection, and that they might be involved in postnatal ZIKV-associated alterations [29]. Infected astrocytes exhibit programmed cell death activation and a mild release of pro-inflammatory cytokines and chemokines [30], [31]. In our previous work, we also described other interrelated cytopathic mechanisms including glial reactivity, mitochondrial failure, oxidative stress and DNA damage in ZIKV-infected astrocytes [19]. Copper has been associated with these alterations in other diseases affecting the CNS [15], [32], [33], and viral infections have been shown to modulate copper in the organism [23], [24], [34]. Here, we investigated the involvement of impaired copper metabolism in the detrimental effects of ZIKV-infection in astrocytes.

Copper has well known antimicrobial properties and it might play a role in the immune system function, since serum copper levels are increased in inflammatory conditions including infection [35]. However, the role of copper in viral infections has been addressed by few previous studies. In cultures of lung cells infected with influenza A virus, the addition or chelation of copper caused a slight reduction in virus replication, and CTR1 knockdown significantly reduced viral RNA synthesis 24h after infection [22]. Miyamoto et al. (1998) showed that copper chelators inhibited apoptosis and release of viral particles in alveolar cells infected with Influenza A virus. These results indicate that Influenza A virus might use copper to promote its replication. However, this mechanism has not yet been investigated for viruses of the genus *Flavivirus*, and in our study copper chelation did not reduce the percentage of ZIKV-infected cells nor the viral load. As for other *Flaviviridae* virus infections, copper levels were found to be increased in the blood and liver of patients infected with Hepatitis C virus (HCV), contributing to liver damage [23]. In both acute and chronic hepatitis, an increase in copper levels can be observed [37], [38], which is related to liver damage caused by oxidative stress [39]. In Vero cells, treatment with copper chloride and Cu(II)-Imidazole reduced infection by the dengue Virus [40], [41]. On the other hand, treatment with copper sulfate increased West Nile Virus plaque formation in chicken embryo fibroblasts [42]. Considering these studies, there is still conflicting data on the function of copper during viral infection and the outcomes of copper modulation in these conditions should be explored further.

In this work, we demonstrated that ZIKV infection causes a series of alterations on copper homeostasis in hiPSC-derived astrocytes, which include the RNA expression downregulation of one of the copper secretion transporters, ATP7B, as well as the upregulation of copper chaperone ATOX1 and a tendency to the upregulation of chaperone COX17. Since these proteins are responsible for targeting copper to organelles or vesicles, as shown in Fig 1a, these changes might indicate the activation of mechanisms aiming copper redistribution, which normally occurs when copper levels are altered. Reduced protein levels of CTR1, which was not caused by downregulation of its mRNA expression, indicates that this protein might be degraded during infection, which is a well-known response against copper overload [43]. Besides copper modulation, ATOX1 is also considered an antioxidant protein, and its transcriptional regulation in response to oxidative stress has already been described in neural damage models [44], [45]. Therefore, the induction of oxidative stress during ZIKV infection in astrocytes may also be responsible for the modulation of ATOX1. We also observed a tendency to COMMD1 upregulation, a multifunctional protein responsible for regulating the stability and localization of ATP7A and ATP7B [46]. COMMD1 also regulates the proteasomal degradation of NF-κB, which is activated as part of the innate immune response against viral infection [47]. In summary, the modulation of copper homeostasis during ZIKV infection in astrocytes could be linked to the activation of the oxidative stress response and innate immune response. Still, more research is needed to confirm the connection between these mechanisms.

In our previous study, we observed that oxidative stress contributes to cell damage during ZIKV infection in astrocytes [19], and we wanted to understand if ZIKV-induced copper modulation is involved in ROS-induced cytotoxicity. Although some of the alterations in copper proteins could indicate an increase in intracellular copper, treatment with CuCl_2_ did not induce further oxidative stress or cell death. This suggests that copper regulatory mechanisms might still be partially functional and able to counteract additional extracellular copper. On the other hand, BCS treatment reduced ROS levels in the mitochondria and cytoplasm and improved cell viability in astrocytes infected with ZIKV. It is known that copper excess can trigger oxidative stress through the formation of ROS in a Fenton-like reaction [48], and oxidative damage through copper overload has already been described in cultured astrocytes and neurons [49], [50]. However, this mechanism had never been described in the context of a viral infection, although some studies hypothesized a relationship between high serum copper levels and oxidative damage in patients with Hepatitis B and C infection [39], [51], [52]. Our data indicate that copper overload might be involved in the oxidative damage and cell death caused by ZIKV infection in astrocytes, but does not seem to involve alterations in the life cycle of the virus, since viral replication or infectivity were not affected by copper chelation or addition. Finally, copper imbalance could also be involved in other cytotoxic effects of ZIKV that are beyond the scope of this work, including disturbances in mitochondrial respiration, DNA damage, proliferation defects and neuroinflammation [15], [32], [53]–[55].

This study focused on astrocytes due to their crucial function in CNS physiology and development and also in the regulation of copper in the brain. The precise control of copper is fundamental in other tissues like the blood-brain barrier and the placenta, which regulate the availability of copper and other compounds to the brain and fetus, respectively. Since these tissues are permissive to ZIKV, it would be important to understand if copper metabolism in these structures is also modified during infection. At the same time, it is also important to consider how ZIKV may be affecting other essential elements such as zinc or iron, which are closely related to copper in terms of biodistribution and function. Notably, zinc and copper share several regulatory proteins, and homeostasis of both biometals is essential for brain physiology [56]. Zinc is considered an antioxidant agent for its ability to induce the expression of metal-binding molecules such as metallothioneins and glutathione [57], and it has a protective role in copper-induced damage [58], [59]. Therefore, it would be relevant to investigate both copper and zinc levels during ZIKV infection. At the same time, zinc presents as a possible therapeutic candidate for the changes in copper homeostasis observed in this work.

## Conclusions

With the increasingly connectedness of the world, disease outbreaks like Zika and COVID-19 are likely to be more frequent and possibly more harmful in the future. It has become clear that understanding the mechanisms of action of the viruses is a very important step towards a fast and effective treatment of the infection. Our work addressed a novel perspective of ZIKV pathology concerning the dysregulation of copper homeostasis. Although some studies have already evaluated the involvement of copper in the pathology of other viral infections, this work is the first to study changes caused by a viral infection in the homeostasis of copper. Given the evidence shown here that copper regulation disturbance is associated with the deleterious effects of ZIKV infection in astrocytes, we highlight the importance of studying the homeostasis of copper and other metals in viral infections and in other pathologies in which this perspective has not been explored yet. Finally, it would be interesting to assess whether changes in copper homeostasis during ZIKV infection occur in other cell types and tissues, or whether it is a specific event during astrocytes infection.

## Acknowledgements

This work was sponsored by *Fundação de Amparo à Pesquisa do Estado do Rio de Janeiro* (FAPERJ), *Conselho Nacional de Desenvolvimento Científico e Tecnológico* (CNPq), Coordenação de Aperfeiçoamento de Pessoal de Nível Superior (CAPES), *Instituto Nacional de Neurociência Translacional (INNT)* and *Banco Nacional de Desenvolvimento* (BNDES), in addition to intramural grants from D’Or Institute for Research and Education. The authors would like to thank Ismael Carlos da Silva Gomes, Gabriela Lopes Vitória and Severino Galdino from D’Or Institute for Research and Education for technical support, and Karina Karmirian, Marília Zaluar P. Guimarães, Sylvie DeValle, Pablo Trindade, Daniel Cadilhe and Daniel Furtado for scientific discussions and input.

## Data availability

The datasets generated and analyzed during the current study are available from the corresponding author on reasonable request.

