## Supplementary Material for "Copper imbalance linked to oxidative stress and cell death during Zika virus infection in human astrocytes"

### SUPPLEMENTARY MATERIALS

**Supplementary Table 1. Primers used for ZIKV entry receptors detection and gene expression analysis**

| Gene | Primer 5' → 3' (*Forward*) | Primer 5' → 3' (*Reverse*) |
|---|---|---|
| AXL | CCGTGGACCTACTCTGGCT | CCTTGGCGTTATGGGCTTC |
| Gas6 | CTCGTGCAGCCTATAAACCCT | TCCTCGTGTTCACTTTCACCG |
| Tyro3 | CGGTAGAAGGTGTGCCATTTT | CGATCTTCGTAGTTCCTCTCCAC |
| TIM-1 | TGGCAGATTCTGTAGCTGGTT | AGAGAACATGAGCCTCTATTCCA |
| TIM-4 | TCCGCACTGATGGAATGAGG | CTTTCACTGGGGTTTAAGATGGT |
| GAPDH | CCATCTTCCAGGAGCGAGATC | TGAAGACGCCAGTGGACTC |
| HPRT1 | CGTCGTGATTAGTGATGATGAACC | AGAGGGCTACAATGTGATGGC |
| CTR1 | GGGGATGAGCTATATGGACTCC | TCACCAAACCGGAAAACAGTAG |
| ATP7A | TGACCCTAAACTACAGACTCCAA | CGCCGTAACAGTCAGAAACAA |
| ATP7B | GCCAGCATTGCAGAAGGAAAG | TGATAAGTGATGACGGCCTCT |
| ATOX1 | GTGCTGAAGCTGTCTCTCGG | GCCCAAGGTAGGAAACAGTCTTT |
| CCS | AGCGGCCAGTTGCAGAATC | CGTCAATAGTTCCCTCGATGAG |
| COX17 | TGCGTGTATCATCGAGAAAGGA | GCCTCAATTAGATGTCCACAGTG |
| COMMD1 | CTGTTGCCATTATAGAGCTGGAA | GCGTCTTCAGAATTTGGTTGACT |

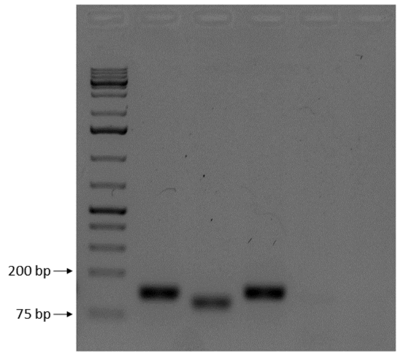


**Supplementary Figure 1. Expression of ZIKV entry molecules in human iPSC-derived astrocytes.** RNA was extracted from control astrocytes for analyzing the expression of ZIKV entry molecules AXL, Gas6, Tyro3, TIM-1 and TIM-4.

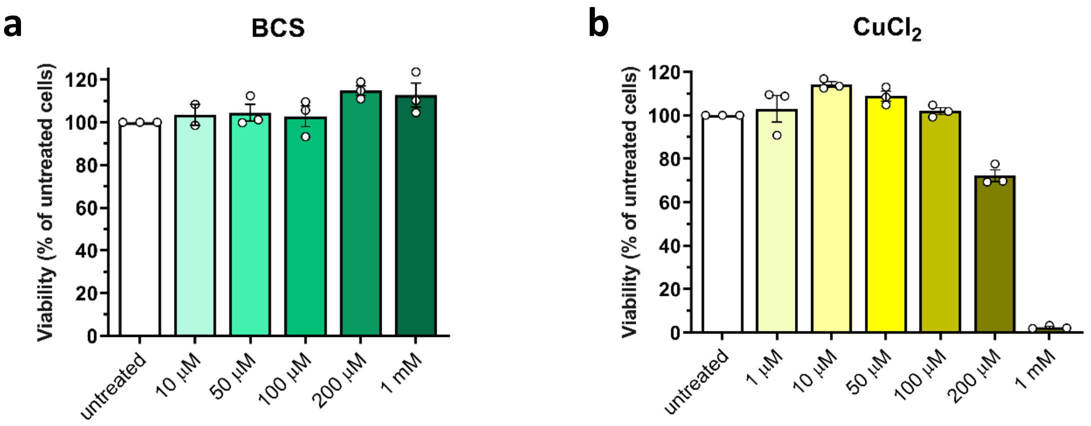


**Supplementary Figure 2. BCS and $CuCl_2$ toxicity analysis on astrocytes.** Astrocytes were treated with BCS (a) or $CuCl_2$ (b) and 96 hours later toxicity was assessed using the neutral red assay. Relative cell viability was calculated considering the untreated group (CTRL) as 100% viable.

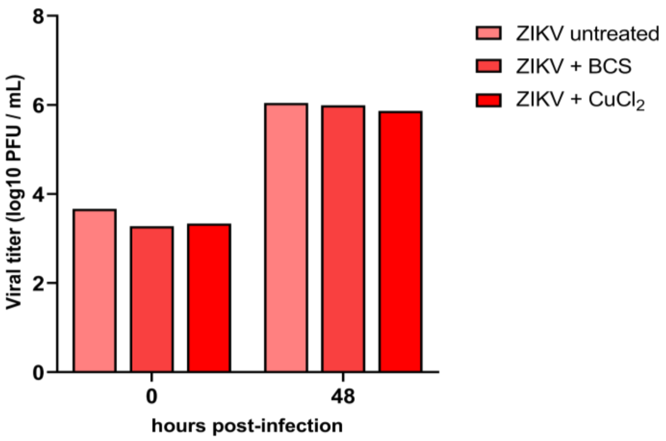


**Supplementary Figure 3. BCS and $CuCl_2$ treatment did not change ZIKV replication in astrocytes.** Astrocytes were infected with ZIKV and treated with BCS or $CuCl_2$, and conditioned medium was collected 0 and 48 h.p.i. for quantification of infectious virus particles by plaque assay. Results were expressed as $log_{10}$ PFU/mL. n=1
